# Strain diversity drives heterogeneous responses to tuberculosis combination therapy

**DOI:** 10.1101/2025.10.06.680645

**Authors:** Michelle H. Yoon, Peter H. Culviner, Mariana Pereira Moraes, Hidetomi Nitta, Nguyen Thuy Thuong Thuong, Sarah M. Fortune, Bree B. Aldridge

**Affiliations:** Department of Molecular Biology and Microbiology, Tufts University School of Medicine, Boston, MA, USA; Department of Microbial Pathogenesis, Yale School of Medicine, New Haven, CT, USA; Department of Immunology and Infectious Diseases, Harvard T.H. Chan School of Public Health, Boston, MA, USA; State University of New York Upstate Medical University, Syracuse, NY, USA; Oxford University Clinical Research Unit, Ho Chi Minh City, Vietnam; Centre for Tropical Medicine and Global Health, Nuffield Department of Medicine, University of Oxford, Oxford, UK; Stuart B. Levy Center for Integrated Management of Antimicrobial Resistance, Tufts University, Boston, MA, USA; Department of Biomedical Engineering, Tufts University School of Engineering, Medford, MA, USA

## Abstract

**Background:** Strain diversity in *Mycobacterium tuberculosis* (Mtb) underlies distinct clinical presentations and outcomes, but the range of drug susceptibility phenotypes among clinical isolates is poorly understood. We aimed to identify drug response patterns in phylogenetically diverse clinical isolates to combination treatment.

**Methods:** Out of 641 drug-sensitive clinical isolates, we selected 13 strains that capture local and global phylogenetic diversity and included Erdman ATCC-35801 as a reference. We selected ten antibiotics with diverse mechanisms of action to study phenotypic responses to combination therapy. We treated each strain with 10 single drugs, 45 drug pairs, and 20 three-way combinations in standard and cholesterol-rich media. To compare combination treatment responses across strains and conditions that have varying doubling times, we computed normalized growth rate inhibition metrics (GR_max_).

**Findings:** Mtb clinical strains displayed a broad range of drug response phenotypes across the 65 drug combinations and two metabolic conditions tested. The most effective drug pairs (based on potency and synergy) varied both by strain and metabolic condition. Within our 14-strain panel, strains that were less sensitive to single drugs were also less sensitive to combination treatment, with very few exceptions. For all drug combinations tested, the magnitude of GR_max_ variation across all strains was driven primarily by variation among genetically related strains, rather than between genetically distant strain groups.

**Interpretation:** Preclinical studies should reflect the diversity of Mtb clinical strains; our data suggest that selecting strains based on the range of drug response phenotypes displayed, rather than by genetic diversity alone, may better account for the effects of strain variation. Our findings also support the understanding that constituent drug pairs of high-order combinations target metabolically heterogeneous Mtb. Selection of these pairs should likely involve multiple factors including the infecting strain, metabolic niche, and drug response metrics.

**Funding:** Gates Foundation INV-027276; NIH P01AI143575&1F32AI174653; Wellcome 206724/Z/17/Z

## Introduction

Tuberculosis (TB) remains challenging to treat, in part because a subset of patients requires extended treatment. Hard-to-treat disease may have several origins: lesions associated with severe pathologies impede drug penetration and drug susceptibility, and genetic differences among infecting organisms—and the phenotypes they mediate—can manifest in heterogeneous disease presentations and responses to drug treatment.^1–4^ Though several studies demonstrate substantial heterogeneity in antibiotic susceptibility, transmission potential, and treatment outcomes among newly emerging *Mycobacterium tuberculosis* (Mtb) clinical strains,^5^ bacterial determinants (beyond canonical resistance mutations) associated with unfavorable clinical outcomes have only recently been characterized.^3^ Recent studies show that strain-to-strain variation in response to antibiotic and metabolic stress and modest differences in MIC values below clinical breakpoints underlie distinct clinical phenotypes such as treatment failure and cavitary disease.^3,6^ However, despite growing evidence that subtle differences in antibiotic susceptibility can lead to divergent clinical outcomes, preclinical treatment design relies on data from commonly studied Mtb strains (Erdman, H37Rv, and HN878) and does not consider the range of drug susceptibility patterns of clinical isolates.^7–10^

Though phenotypic responses to several first- and second-line antibiotics are well characterized across different Mtb phylogenetic lineages,^11–13^ TB treatment relies on combination therapy, and clinical outcomes likely reflect strain-dependent differences in drug combination efficacy.^14^ Yet, we lack a systematic evaluation and understanding of combination treatment responses across diverse clinical isolates. Here, we provide an analysis of drug responses to 65 unique drug combinations across a phylogenetically diverse panel of Mtb clinical strains in two *in vitro* conditions that model different cell states. Our data allows us to answer several fundamental questions about the impacts of strain variation on combination treatment outcomes: How variable in drug susceptibility are the clinical isolates to single drugs and drug combinations? Do genetically related isolates exhibit similar drug susceptibility patterns? Are certain isolates more tolerant to certain combinations, and if so, can we identify any drug-dependent or strain-dependent patterns? Finally, can we reasonably predict how susceptible a given isolate is to combination treatment based on its susceptibility to single drugs - that is, can we screen problematic isolates based on single drug susceptibility? The variation we highlight across strains and drug combination treatments, in different metabolic conditions, may aid future combination treatment optimization in accounting for pathogen diversity, with implications for de-risking, revamping, and accelerating drug regimen design.

## Methods

### Antimicrobials, dispensing, and drug combination design

A complete list of antibiotics used in this study is provided in Supplementary Table 1. 50% inhibitory concentrations (IC_50_) were calculated for all antibiotics in two different *in vitro* models with different carbon sources: standard (sugar-rich) and cholesterol (lipid-rich). Acclimation procedures are detailed in Appendix 1. For drug combinations, equipotent mixtures of antibiotics were created by aligning the IC_50_ value of each constituent drug to the same dose level.^15^ Initial IC_50_ measurements were made using 14 doses with 2-fold spacing, whereas subsequent experiments used a 1.5-fold dose resolution across 10 doses.

### Strains and culturing

All strains used in this study were prepared and cultured in a 7H9 broth supplemented with 0.05% Tween 80, 0.2% glycerol, and 10% Middlebrook OADC (referred to as standard medium in the main text) and stored in -80°C until use. Frozen 1ml stocks of Mtb were added to 9ml of standard medium and grown in a 37°C shaking incubator to mid-log phase (optical density at 600nm, or OD_600_, between 0.4 and 0.8).

### Drug treatment assays

Dose response assays for drug combinations were designed using the DiaMOND (diagonal measurement of *n*-way drug interactions) method, which optimizes traditional checkerboard assays to measure equipotent mixtures of drugs.^16^ This allows efficient measurements of drug interactions in a large number of drug combinations that would otherwise be impractical with traditional methods.

50µl of Mtb grown in either standard or cholesterol medium was back-diluted to an OD_600_ of 0.05, added to drug-treated 384-well plates, and incubated at 37°C in humidified bags to prevent evaporation. Edge wells (top and bottom rows, left- and right-most columns) contained only media and were excluded from analysis. OD_600_ was measured at two time points for each strain using a Synergy Neo2 Hybrid Multi-Mode Reader. For both standard and cholesterol conditions, OD_600_ was measured immediately following inoculation (day 0) and at the terminal time point, which corresponds to 5 days post-inoculation (Day 5) for standard and 12 days post-inoculation (day 12) for cholesterol. These time points were identified as being predictive of treatment outcomes *in vivo.*^8^

### Data processing and DiaMOND metric calculation

OD_600_ data were processed using custom MATLAB analysis scripts. Raw data were first derandomized and then background-subtracted using the median OD_600_ value of the media-only edge wells. Drug-treated wells were then normalized to the mean OD_600_ value of the untreated controls on the same plate. To generate dose response curves ranging from 0 (no growth inhibition) to 1 (complete growth inhibition), normalized OD_600_ values were subtracted from 1. Each resulting dose-response curve (single drug or drug combination) was fit to a three-parameter Hill function, and the resulting Hill curve parameters were used to calculate inhibitory concentrations (ICs). Calculation of growth rate inhibition (GR) and drug interaction metrics are detailed in Appendix 1.

### Phylogenetic tree construction

A core genome phylogenetic tree was constructed by downloading complete genomes for the relevant bacterial strains from NCBI. The Erdman ATCC 35801 strain was used as the reference genome. Paired-end FASTQ files for each strain were aligned to the reference genome, and variants were identified for each strain using the ‘bcftools’ package in Python. Core genome alignments were generated, and evolutionary relationships between each strain and the reference genome were summarized using ‘parsnp’ and ‘harvesttools’ packages. A maximum-likelihood phylogenetic tree was then constructed using the ‘IQ-TREE’ package, which evaluates multiple nucleotide substitution models and selects the best-fitting model. Scripts were written in Python 3 and Bash, and the final phylogenetic tree (Figure 1A) was visualized using the ‘tree’ package in R.

**Figure 1.**
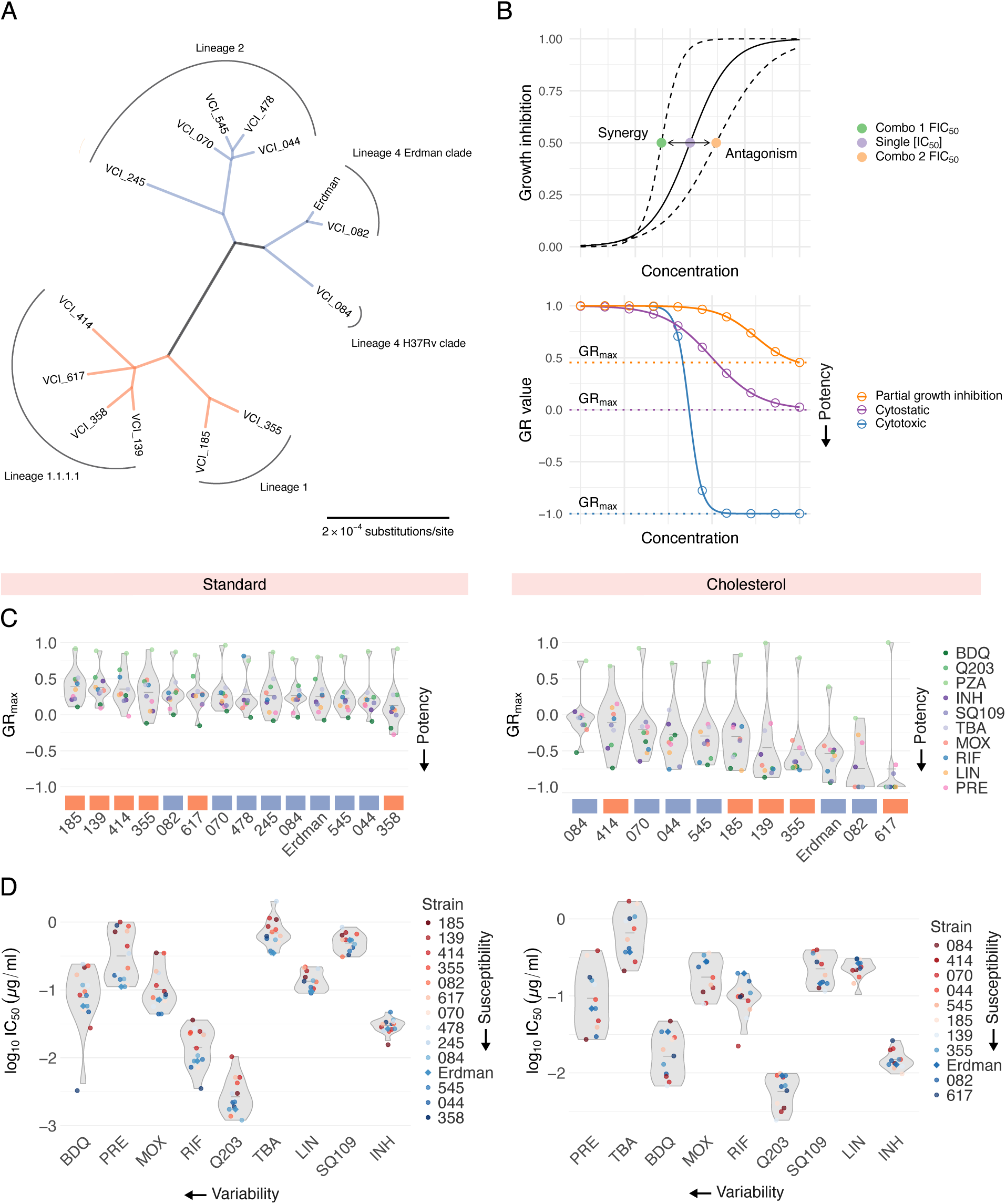
Susceptibility profiles of phylogenetically diverse Mtb clinical isolates to ten antibiotics. A: Unrooted maximum-likelihood phylogenetic tree of the 14 Mtb strains studied in this work, as determined by core gene alignments. Scale bar represents substitutions per nucleotide. Colors indicate strain groups defined for the interpretation of this work. B: Schematics of drug interaction (top) and potency (bottom) metrics computed via DiaMOND. (Top) The solid-line dose response curve represents the inhibitory effect of a single drug. Shifts in the left and right directions indicate synergy and antagonism, respectively, when combined with another drug. (Bottom) Dose response curves indicate a cytotoxic response (blue), cytostatic response (purple), and partial growth inhibition (orange). C: Drug susceptibility profiles of 14 Mtb strains across ten antibiotic treatments in standard (left) and eleven strains in cholesterol (right). Strains are ordered from least drug-susceptible to most drug-susceptible (left to right, respectively). Lineage identity of each strain is represented by orange (Lineage 1) and blue (Lineage 2 and 4) boxes. Each point represents the median GR_max_ value across six biological replicates in standard and three biological replicates in cholesterol. D: Half-maximal inhibitory concentrations (IC_50_) of nine antibiotic treatments across 14 strains in standard (left) and 11 strains in cholesterol (right). PZA is excluded due to inactivity as a single agent. Strains are colored from least drug-susceptible (red) to most drug-susceptible (blue). Antibiotics are ordered from most variable (highest coefficient of variation, CV) to least variable (lowest CV), from left to right, respectively. Each point represents the median log_10_IC_50_ value across six biological replicates in standard and three biological replicates in cholesterol.

### Statistics and data reproducibility

All experiments included in this study were performed in three biological replicates excluding single drug dose response experiments in standard (Figure 1), where six biological replicates were used instead. We observed greater variability for certain drugs in standard and aimed to increase precision with the use of additional biological replicates. For each strain and drug treatment, the median log_10_IC_50_ (for single drugs), log_2_FIC_50_ (for drug combinations), and GR_max_ (for drug combinations) value across three biological replicates (for drug combinations and single drugs in cholesterol) and six biological replicates (for single drugs in standard) were measured and represented in the figures.

## Results

Clinical isolates were obtained from TB patients with diverse clinical presentations including lung cavitation and treatment failure, as detailed in Stanley et al., 2024.^3^ We selected 13 strains that capture local and global phylogenetic diversity, including two strains from subclades of L1.2 (185 and 355), two strains from the locally expanded L1.1.1.1 (139 and 358), two outgroup strains in L1.1.1.1 (414 and 617), a strain each from the L4.1 Erdman clade and L4.4 H37Rv clade (082 and 084, respectively), a strain from the proto-Beijing L2.1 (245) and four strains from the expanded L2.2 lineage (044, 070, 478, 545) (Figure 1A). We selected ten antibiotics that represent a broad range of mechanisms of action (Table 1) and included Erdman ATCC 35801, commonly used in systematic drug combination studies,^7,8^ as a reference. Based on the phylogenetic tree of the 14 strains, we found that the strains bifurcated into two major groups: Lineage 1 (Figure 1A, orange) and Lineages 2 and 4 (Figure 1A, blue). We manually classified the strains into these two groups for ease of interpretation.

Mtb occupies metabolically distinct niches *in vivo*.^2^ Cholesterol is a key carbon source in lipid-rich environments within granulomas,^17,18^ and studies demonstrate substantial heterogeneity in drug response between Mtb acclimated to different carbon sources.^2,19,20^ Both glycerol- (called standard here) and lipid-based *in vitro* models (including cholesterol) have been validated as predictive of treatment outcomes *in vivo*.^7,8^ To account for condition-dependent variation in drug response phenotypes, we conducted all drug combination assays in standard conditions and in a medium with cholesterol as the sole carbon source (called cholesterol here). We note that a subset of strains—245, 358, and 478—arrested growth in cholesterol and were excluded from analysis in this condition.

We observed substantial variation in doubling times across clinical isolates in both standard (14 to 21 hours) and cholesterol (51 to 104 hours) conditions. To compare drug and drug combination efficacy across different strains and conditions, we used GR metrics, which normalize drug treatment response to the untreated growth rate of each strain. Specifically, we used GR_max_, the maximum growth rate inhibition, to compare drug and drug combination responses across 14 Mtb strains and two *in vitro* conditions. We also computed commonly used drug response metrics such as inhibitory concentrations (ICs) and drug interaction scores (fractional inhibitory concentrations, FICs, Figure 1B, top). A subset of drugs failed to achieve at least 90% growth inhibition in several strains; we therefore used IC_50_ values to interpret phenotypic responses to single drugs and to compute drug interaction scores at the IC_50_ (FIC_50_). Negative, zero, and positive log_2_FIC_50_ values denote synergy, additivity, and antagonism, respectively.

To characterize drug response patterns across our panel of 14 Mtb strains, we treated each strain with ten antibiotics in standard and cholesterol conditions. For each drug treatment, we generated dose response curves from which we calculated GR_max_ and IC_50_ values, resulting in 280 metrics in standard and 220 metrics in cholesterol. In both conditions, we observed strain-to-strain differences in GR_max_ for all ten drugs tested (Figure 1C). For select drugs such as bedaquiline, pretomanid, and moxifloxacin in standard, and bedaquiline, pretomanid, and TBA-3731 in cholesterol, IC_50_ values varied by nearly 10-fold between strains (Figure 1D). We observed that most clinical isolates are more drug tolerant than Erdman in both standard and cholesterol (Figure 1C). Notably, bedaquiline and pretomanid displayed the greatest variability in IC_50_ values across all strains in both conditions (Figure 1D), suggesting that despite their use in new regimens, their efficacy may be inconsistent across clinical strains.

We next asked whether treating Mtb with drug pairs improves treatment efficacy relative to single drugs, based on changes in GR_max_ values. To quantify GR_max_ improvement, we calculated the difference between the GR_max_ of a drug pair and the GR_max_ of its constituent singles, with negative values indicating improvement. Across all strains in standard medium, the addition of bedaquiline or pretomanid to another drug improved the GR_max_ of the other drug, with improvement observed in 216/224 (96.5%) drug pairs containing either bedaquiline or pretomanid (Figure 2A, left). Given that bedaquiline and pretomanid are the two most potent single drugs in standard medium, we hypothesized that combining any drug with another drug that is less potent than itself will invariably improve the GR_max_ of the less potent drug. We found that this was generally the case, but not to the same extent as bedaquiline or pretomanid. For instance, SQ109, linezolid, and isoniazid also improved the GR_max_ of drugs less potent than itself in a majority of drug pairs (84/98 drug pairs or 85.7%, 69/84 drug pairs or 82.1%, 67/70 or 95.7%, respectively), but they improved the GR_max_ of more potent drugs (such as bedaquiline and pretomanid) in only a subset of strains (Supplementary Figure 1, top). However, we did not observe this pattern in other drugs such as moxifloxacin or rifampicin—in these drugs, GR_max_ improvement was less predictable (Supplementary Figure 1, top). These findings suggest that (i) in some cases, GR_max_ improvement of drug pairs can be predicted based on the potency of the individuals drugs being combined and (ii) bedaquiline and pretomanid may have intrinsic properties that allow them to optimize drug combination efficacy in diverse strain backgrounds beyond what would be expected based on potency alone. In cholesterol, however, no single drug improved GR_max_ across all combinations and strains (Figure 2A, right; Supplementary Figure 1, bottom). Instead, the magnitude of GR_max_ improvement of any given drug pair varied by strain, suggesting that combination efficacy in cholesterol depends more on strain-specific susceptibility patterns to drug pairs than by the intrinsic properties of individual drugs.

**Figure 2.**
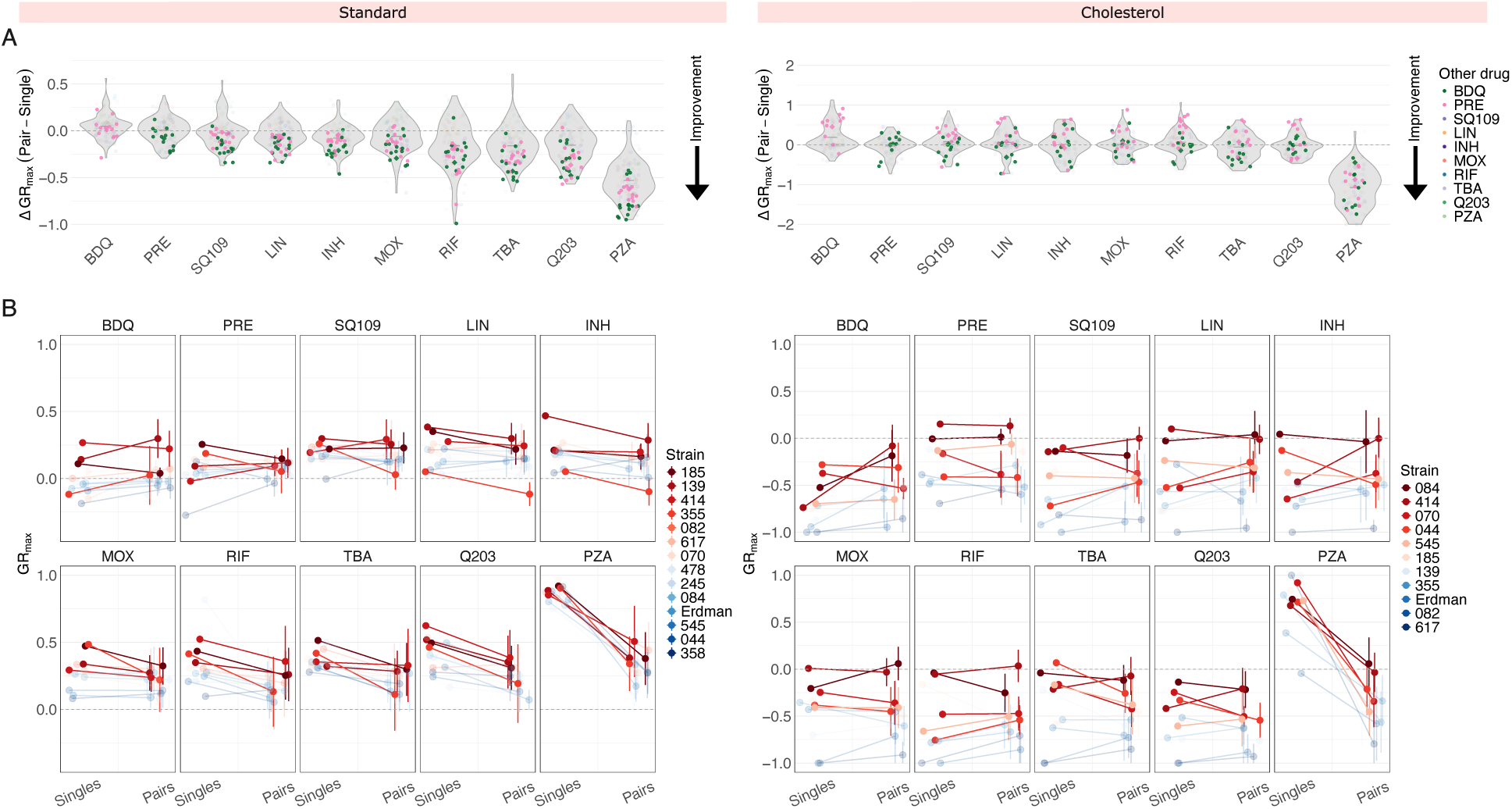
Comparison of GR_max_ values between different drug modalities. A: Change in GR_max_ between single drug and pairwise treatments in standard (left) and cholesterol (right) conditions. Each point denotes a ΔGR_max_ value of a drug pair for an individual isolate and represents the median GR_max_ value across three biological replicates. Drug pairs containing bedaquiline and pretomanid are highlighted in green and pink, respectively. A negative ΔGR_max_ value indicates an improvement in GR_max_ when a single drug is combined with a second drug (e.g., when BDQ is combined with LIN, or when INH is combined with MOX). B: Comparison of GR_max_ values between single drugs and their nine corresponding drug pairs (Supplementary Table 2) in standard (left) and cholesterol (right) conditions. Each panel represents a single drug on the left side and the mean GR_max_ value of its nine corresponding drug pairs on the right side. Vertical bars represent the standard deviation of the mean GR_max_ value. Each point represents the median GR_max_ value across three biological replicates and corresponds to an individual strain. The four least susceptible strains for each condition are highlighted in red. (Note: opacity adjustments are not reflected in the legend.)

The drug susceptibility profile of a specific Mtb strain is typically defined based on its minimum inhibitory concentration (MIC) values to select single drugs, even though TB patients are treated with drug combinations. We therefore asked whether drug response phenotypes to single drugs provide any predictive insight into strain-specific phenotypic responses to combination treatment. We compared GR_max_ values across three different treatment modalities—single drugs (10 drugs), drug pairs (45 pairs), and three-way combinations (20 combinations)—for each strain (Figure 2B; Supplementary Figure 2; Supplementary Table 2; Supplementary Table 3). This comparison addressed two related questions: (1) within our 14-strain subset, are strains that are more difficult to treat with single drugs also more difficult to treat with drug combinations, and (2) between different treatment modalities, does GR_max_ change in a strain-dependent manner, or does it follow a generalizable trend? In both standard and cholesterol, we found that strains that are less sensitive to single drugs are also less sensitive to drug pairs and three-way combinations on average (Figure 2B; Supplementary Figure 2), as these strains (represented in red) occupy the upper range in GR_max_ across nearly all drugs in the three treatment modalities tested (Figure 2B; Supplementary Figure 2). We conclude that the stratification of strains by single-drug susceptibility is largely preserved between treatment modalities. An exception was isolate 355 in standard medium; although more tolerant to single drugs than most other strains, its GR_max_ was greatly improved by treatment with drug pairs, especially those containing SQ109, moxifloxacin, rifampin, and TBA-3731 (Figure 2B, left).

We then evaluated whether three-way combinations provide further improvement in treatment efficacy over drug pairs. Given that only six drugs were tested in three-way combinations, as opposed to ten drugs in drug pairs, comparisons were limited to shared compounds. In both standard and cholesterol, there was no improvement in GR_max_ in three-way combinations over drug pairs, suggesting that high-order combinations do not provide additional improvement over drug pairs in the conditions tested (Supplementary Figure 2).

Drug potency and strain variability metrics of (A) 45 drug pairs and (B) 20 three-way combinations tested in standard and cholesterol. For each condition: (top) violin plots represent the GR_max_ range across all clinical strains tested. The four least drug-susceptible strains and the four most drug-susceptible strains (Fig. 1D) are highlighted in red and blue, respectively. All other strains are represented in grey. Each point represents the median GR_max_ value across three biological replicates. (Middle) Each bar represents the coefficient of variation (CV) of GR_max_ values across 14 strains in standard and 11 strains in cholesterol for each drug pair. (Bottom) Each stacked bar represents the variance decomposition of the GR_max_ values across 14 strains in standard and 11 strains in cholesterol for each drug pair. The green bar represents the percent total variance attributable to variance between lineage 1 and lineages 2 and 4; the orange bar represents the percent total variance attributable to variance within lineage 1; the blue bar represents the percent total variance attributable to variance within lineages 2 and 4.

To improve the resolution of strain-specific phenotypic responses to drug combination treatment, we decomposed averaged drug combination data into all individual drug pairs (Figure 3A) and three-way combinations (Figure 3B) tested. This allowed us to ask several questions: (1) are hard-to-treat strains (defined by low susceptibility to single drugs) more tolerant to all drug combinations, or do specific combinations sensitize them, and (2) do some drug combinations display larger variability in GR_max_ than others, and if so, do certain strains drive this variability? We found that in both standard and cholesterol conditions, hard-to-treat strains are more tolerant to nearly all drug pairs and three-way combinations tested (Figure 3A and 3B, violin plots), suggesting that strains that are tolerant to single drugs are expected to be more difficult to treat with drug combinations. Regardless, a small subset of drug combinations was effective against select hard-to-treat strains. For example, of all 45 drug pairs and 20 three-way combinations tested, pretomanid+rifampin (PRE+RIF), pretomanid+telacebec (PRE+Q203), and isoniazid+pretomanid+rifampin (INH+PRE+RIF) were the only drug combinations that induced a cytotoxic effect in isolate 139 in standard (Figure 3A and 3B, violin plots for standard). Similarly, bedaquiline+isoniazid (BDQ+INH) and bedaquiline+TBA-3731 (BDQ+TBA) also showed increased potency against isolate 185 (Figure 3A, violin plot for standard). These examples highlight the potential of targeted combinations to sensitize strains that may otherwise be tolerant to a broad swath of treatment options.

**Figure 3.**
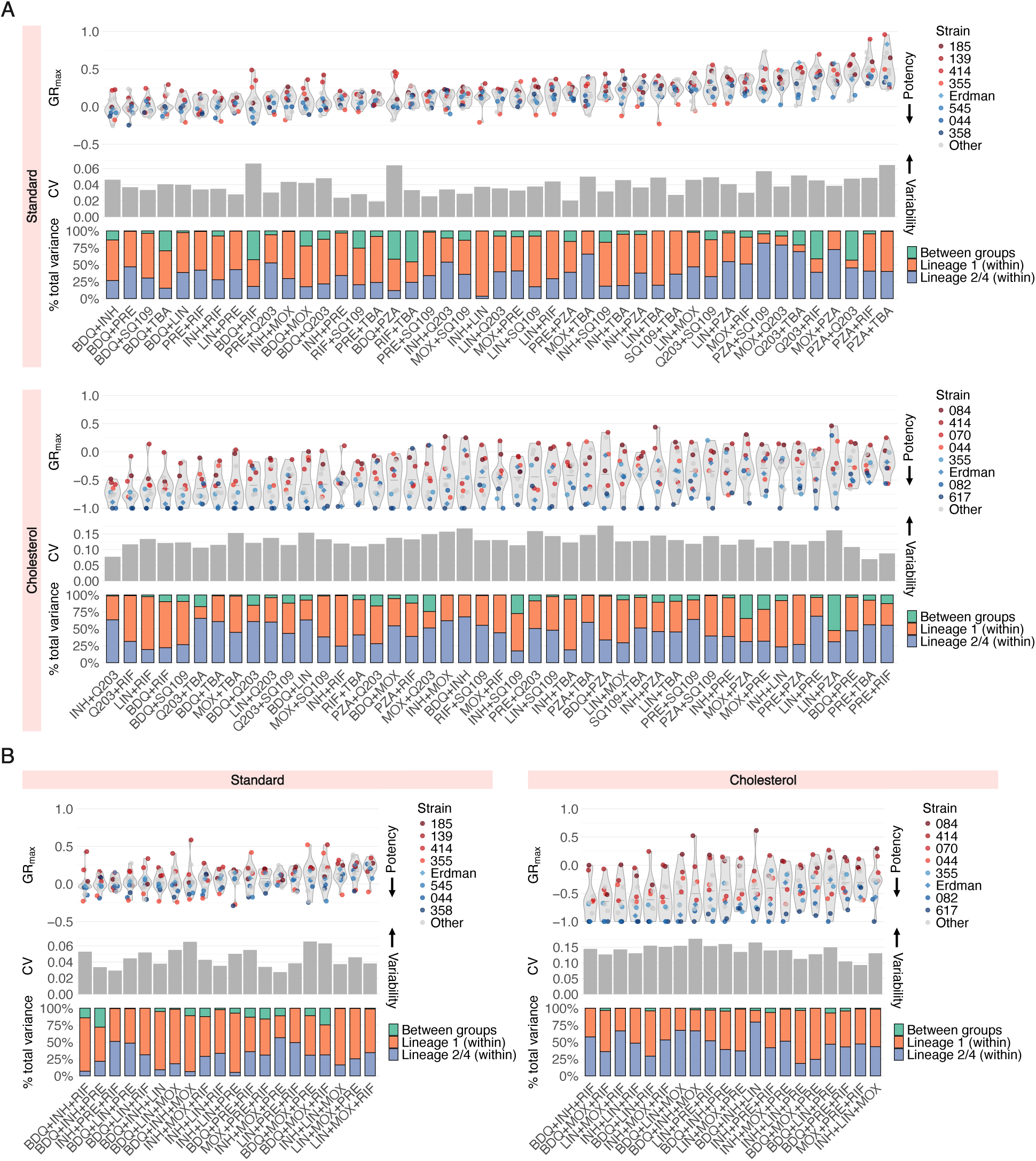
Susceptibility profiles of Mtb clinical isolates to drug pairs and high-order combinations.

Across all drug pairs and three-way combinations, we sought to identify a quantitative relationship between drug combination potency, as determined by GR_max_, and the magnitude of strain-to-strain variability, as determined by the coefficient of variation (CV) (Figure 3A and 3B, gray bar plots). There was no correlation between combination potency and the magnitude of strain-to-strain variability, but a small subset of potent drug combinations—bedaquiline+rifampin (BDQ+RIF), bedaquiline+pyrazinamide (BDQ+PZA), and bedaquiline+isoniazid+moxifloxacin (BDQ_INH+MOX) in standard medium—were more variable than others in the same condition (Figure 3A and 3B, gray bar plots for standard). Notably, we found that strain-to-strain variability in GR_max_ to single drugs (Supplementary Figure 3) is not correlated with variability in GR_max_ to drug pairs (Figure 3A, gray bar plots for standard). That is, although bedaquiline and pretomanid exhibit the most strain-to-strain variability in GR_max_ (Supplementary Figure 3), combining pretomanid with another drug tended to reduce the amount of variability; in contrast, pairing bedaquiline with another drug tended to increase variability (Figure 2B, left). Moreover, we found that three-way combinations do not reduce variability relative to drug pairs (Figure 3A and 3B, gray bar plots; Supplementary Figure 4), suggesting that increasing the number of drugs does not mitigate strain-to-strain differences in single drug response.

We next asked whether specific strains or strain groups drive the variation in GR_max_ observed in each of the drug combinations tested. We partitioned the total variance in GR_max_ for each drug combination into three components: variance within lineage 1 strains, variance within strains in lineages 2 and 4, and variance between the two strain groups (Figure 3A and 3B, stacked bar plots). Strikingly, for all combinations tested, the total variance was driven primarily by variability in GR_max_ within strain groups, rather than between (Figure 3A and 3B, stacked bar plots), suggesting that genetically related strains do not necessarily exhibit similar drug response phenotypes.

Though we demonstrate that three-way combinations do not improve drug potency over drug pairs in any one condition, TB treatment still necessitates high-order combinations since its constituent drug pairs may target different metabolic and physical niches. We sought to identify candidate drug pairs, based on their GR_max_ and FIC_50_ values, that could potentially be effective across diverse strain backgrounds and different *in vitro* conditions. A few drug pairs were both potent and synergistic in standard and cholesterol (Figure 4A, orange), but only a small subset of these pairs—pretomanid+rifampin (PRE+RIF) and rifampin+SQ109 (RIF+SQ109)—were shared between the two conditions (Figure 4A, orange). For all drug pairs, the standard deviation in potency (horizontal error bars) and synergy (vertical error bars) varied greatly, suggesting that drug pairs that satisfy both criteria (potent and synergistic) vary by individual strain. We further demonstrate limited correlation between potency and synergy, as drug pairs that are potent in both conditions (Figure 4B, left) and synergistic in both conditions (Figure 4B, right) vary. Altogether, these data further support our understanding that different drug pairs target metabolically heterogeneous Mtb, and the selection of these pairs relies on a multitude of factors including genetic background, metabolic state, and a consideration of several drug response metrics.

**Figure 4.**
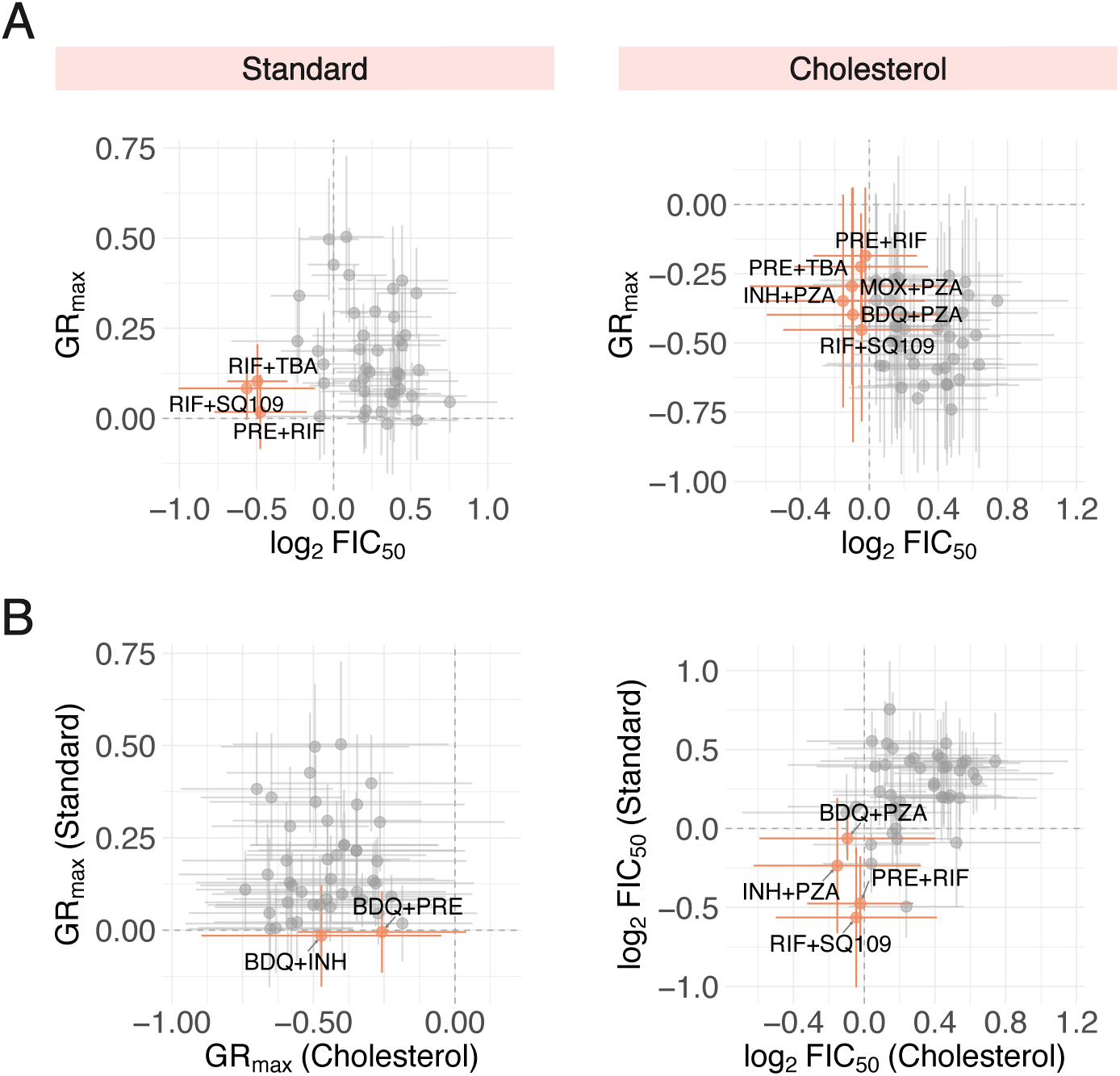
Comparison of drug pairs combination treatment responses across different *in vitro* conditions and dose response metrics. A: Potent (low GR_max_ value) and synergistic (low log_2_FIC_50_ value) drug pairs identified in *in vitro* conditions. (Left) The three drug pairs with the lowest mean GR_max_ and log_2_FIC_50_ values are represented by orange points. All other drug pairs are represented by grey points. Vertical and horizontal bars denote standard deviation of GR_max_ and log_2_FIC_50_ values, respectively. (Right) Drug pairs that are both cytotoxic and synergistic, indicated by negative GR_max_ and log_2_FIC_50_ values, respectively, are represented by orange points. B: (Left) Drug pairs that are cytotoxic (negative GR_max_ value) in both standard and cholesterol conditions are represented by orange points. Vertical and horizontal bars denote the standard deviation of GR_max_ values and standard and cholesterol, respectively. (Right) Drug pairs that are synergistic (negative log_2_FIC_50_ value) in both standard and cholesterol conditions are represented by orange points.

## Discussion

Altogether, we demonstrate substantial heterogeneity in drug response to combination treatment across our 14-strain panel, with certain strains exhibiting attenuated sensitivity to nearly all drugs and drug combinations tested. Importantly, many clinical isolates were less sensitive to combination treatment than commonly studied strains such as Erdman, and treatment efficacy of drug combinations varied greatly by strain, especially those that included drugs that are relevant in the clinic and in preclinical studies. Our findings highlight the importance of studying the effects of drug combination treatments in phenotypically diverse Mtb strains, particularly to define the upper bounds of drug tolerance.

While identifying genetic resistance markers remains essential in understanding the causes of severe disease pathologies and treatment failure, an increasing body of work shows that physiological adaptations such as altered respiration and energy metabolism, metabolic reprogramming, and upregulation of multidrug efflux pumps can broadly reduce drug susceptibility independent of genotype.^21–24^ Our results support the understanding that such mechanisms, not readily identified via genomic analyses alone, may underlie broad-spectrum drug tolerance to a larger extent than expected: strains that are less susceptible to single agents often remain recalcitrant to high-order drug combinations, even when those combinations include drugs with distinct mechanisms of action. That is, increasing the number of drugs in a combination failed to mitigate relative drug tolerance, or reduce the magnitude of strain-to-strain variability in drug response. These findings suggest that high-order combinations alone are insufficient to rescue certain forms of intrinsic or acquired drug tolerance.

Strikingly, strain-to-strain variation in drug response was driven primarily by the susceptibility profiles of individual strains, rather than by lineage identity. Strain phenotypes are likely driven by mutations arising on terminal branches, or other contemporaneous changes that are heritable yet appear randomly distributed across the phylogenetic space due to our limited resolution (n = 14 strains). Additionally, adaptations shaped by various facets of the host environment that modulate drug response may have been selected for during infection and retained *in vitro*. Our results also have implications in designing strain panels for preclinical combination design studies; we propose that selecting strains based on phenotypic, in addition to genetic, diversity may more effectively capture strain diversity. Overall, our findings advocate for a holistic approach to studying the effects of combination therapy—one that probes both the upper and lower bounds of drug susceptibility across phenotypically diverse strains and contextualizes these phenotypes in several host-relevant metabolic conditions.

There are several limitations to our study. First, although our 14-strain panel spans both local and global phylogenetic diversity, it is derived from a single geographic cohort and thus represents a limited fraction of the global Mtb population. Further expanding the strain panel in scope and size may provide new insight into drug response traits underlying heterogeneous treatment outcomes. Second, though our cholesterol model reflects host-relevant carbon sources, it does not fully recapitulate the lesion microenvironment such as immune pressure, hypoxic stress, and nutrient restriction. Additionally, the cholesterol condition alone only represents a fraction of the metabolic diversity of Mtb during host infection. Future combination studies should include multiple host-like *in vitro* conditions to account for this diversity. Finally, a subset of strains arrested growth in cholesterol medium and were thus excluded from further analysis in that condition, limiting direct comparisons across the complete strain panel.

The current first-line regimen and several other drug combinations used in the clinic vary in treatment efficacy among TB patients.^25,26^ To mitigate this variability, it is crucial for key processes in drug combination design and optimization to account for strain-to-strain differences in drug susceptibility. Our study demonstrates that strain variation in drug combination responses may explain some of the differences in drug responses observed in the clinic, which suggests that introducing strain diversity into these preclinical studies has major implications in optimizing and derisking drug regimen design.

### Contributors

MHY, PHC, SMF, and BBA were responsible for conceptualization of the study. MHY and BBA conducted the literature search, developed the figures, and performed formal analysis and writing of the original manuscript. MHY, MPM, and HN were responsible for data curation. MHY, PHC, SMF, and BBA contributed to data interpretation. PHC, NTTT, SMF, and BBA were responsible for funding acquisition and supervision of the study. All authors had access to the study data and contributed to the review and editing of the manuscript.

## Data availability

Whole-genome sequences of all strains used in this study are available through the National Center for Biotechnology Information Sequence Read Archive (SRA) under the accession number PRJNA950969. Accession numbers of individual strains are listed in Appendix 1 (Supplementary Table 4). All equations used to quantify drug response in this study are listed in the Methods section and Appendix 1 (Supplementary Methods). The *in vitro* data used in this study and the scripts required to generate the figures are available on Figshare.^27–29^

## Acknowledgements

This study was funded by the National Institute of Health (P01AI143575 [SMF], 1F32AI174653 [PHC]), the Wellcome Intermediate Fellowship (2026724/Z/17/Z [NTTT]), and the Gates Foundation (INV-027276 [BBA]). The conclusions and opinions expressed in this work are those of the author(s) alone and shall not be attributed to the Gates Foundation. Under the grant conditions of the Foundation, a Creative Commons Attribution 4.0 License has already been assigned to the Author Accepted Manuscript version that might arise from this submission. Please note works submitted as a preprint have not undergone a peer review process.

## Appendix 1

### Supplementary Methods

#### Antibiotics preparation and dispensing

Each antibiotic was diluted in dimethyl sulfoxide (DMSO) to create a stock solution, which was then prepared into single-use aliquots and stored at -20°C until use.

Dispensing errors were avoided with the use of the HP D300e digital dispenser to dispense drugs and drug combinations into 384-well plates, and well positions were randomized to control for plate effects.

#### *In vitro* model acclimation

Mtb were acclimated to cholesterol-supplemented medium as follows. A base medium containing 7H9 powder (4.7g/L), NaCl (100mM), tyloxapol (0.05%), and fatty acid-free BSA (0.5g/L) was prepared and heated to 37°C, and a cholesterol stock solution was prepared by dissolving cholesterol in a 1:1 mixture of ethanol and tyloxapol and heated to 80°C for 30 minutes. This mixture was then added to pre-warmed base medium at a final concentration of 0.2 mM. Mtb were grown in standard medium to mid-log phase and were sub-cultured once prior to acclimation. Once acclimated to cholesterol medium, cultures were sub-cultured once into fresh medium and grown to mid-log phase prior to drug treatment assays.

#### Growth rate inhibition (GR) metrics calculation

GR values were derived using the following equation^30^:

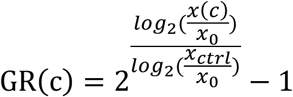

Where x(c) is the OD_600_ value of a drug-treated sample at concentration c at a specific time point, x_ctrl_ is the measurement of the untreated control at the same time point, and x_0_ is the OD_600_ value measured immediately prior to drug exposure (time point 0). GR values quantify the relative growth rate of treated cells compared with untreated cells.

GR was calculated for each drug concentration in the dose–response curve, and a three-parameter Hill function was fit to the data:

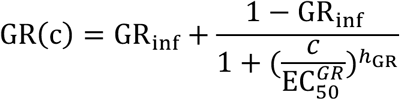

Where GR_()*_ denotes the maximal inhibitory effect, EC^-.^ is the concentration at half-maximal growth rate inhibition, ℎ_01_ is the Hill slope, and c is the concentration of the drug supplied. GR_234_ is the maximum effect of a drug at its highest tested concentration and ranges from -1 to 1. A negative value corresponds to a cytotoxic response, a value of 0 corresponds to a cytostatic response, and a positive value corresponds to partial growth inhibition.

By definition, GRmax values range from -1 (complete cytotoxicity) to 1 (no inhibition). In cholesterol, where the doubling times for x_ctrl_ can exceed 90h for some strains, the denominator in the GR equation, 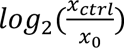 becomes small, making the metric sensitive to minor sources of error such as background noise. Additionally, cholesterol often precipitates over the course of the 12-day incubation step and can increase apparent OD_600_ readings. These effects can occasionally produce computed GR values slightly below -1, which are not biologically possible (as they imply more than 100% loss of cells) and therefore represent numerical artifacts rather than true phenotypes. We therefore bounded all values less than -1 to -1, preserving the intended scale of GR and preventing spurious extreme values from influencing downstream analyses.

#### Drug interaction quantification

For drug combinations, drug interactions were quantified using Loewe additivity as the null model to compute fractional inhibitory concentration (FIC) values. For each drug combination, the expected IC value was calculated by summing the individual IC values of each constituent drug. The FIC value was then computed as the ratio of the experimentally observed IC value of the drug combination to the expected IC value. FIC values were log_2_-transformed to center synergy and antagonism scores at 0. A negative log_2_FIC value indicates synergy, a positive log_2_FIC value indicates antagonism, and a log_2_FIC value at zero indicates additivity. FIC values were calculated at IC_50_ (FIC_50_).

## Supplementary Tables and Figures

**Supplementary Table 1.** Names, abbreviations, and mechanisms of action of the ten antibiotics used in this study. For all ten antibiotics, the commercial name, the abbreviation used in this study, and its mechanism of action are listed in the left-most, middle, and right-most columns, respectively.

| Drug | Abbreviation | Mechanism of action |
| --- | --- | --- |
| Bedaquiline | BDQ | Respiration |
| Telacebec | Q203 | Respiration |
| Pyrazinamide | PZA | Respiration or Multi-effect |
| Isoniazid | INH | Cell wall |
| SQ109 | SQ109 | Cell wall |
| TBA-7371 | TBA | Cell wall |
| Moxifloxacin | MOX | DNA |
| Rifampicin | RIF | RNA |
| Linezolid | LIN | Protein |
| Pretomanid | PRE | Multi-effect |

**Supplementary Table 2.**
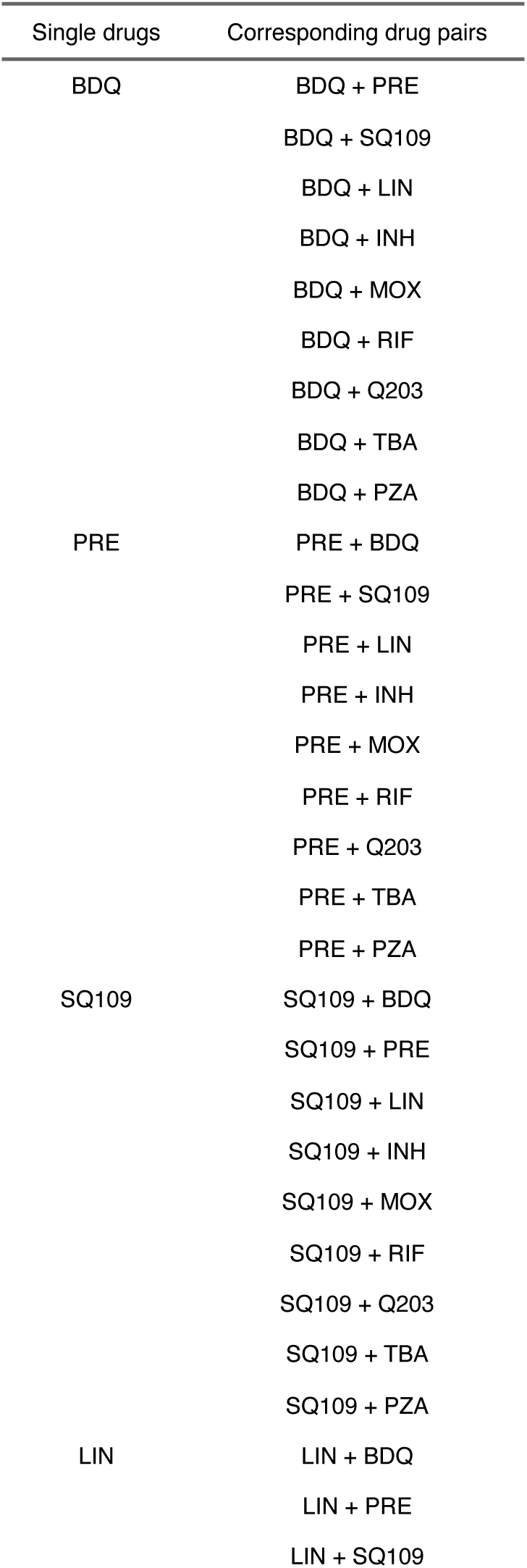
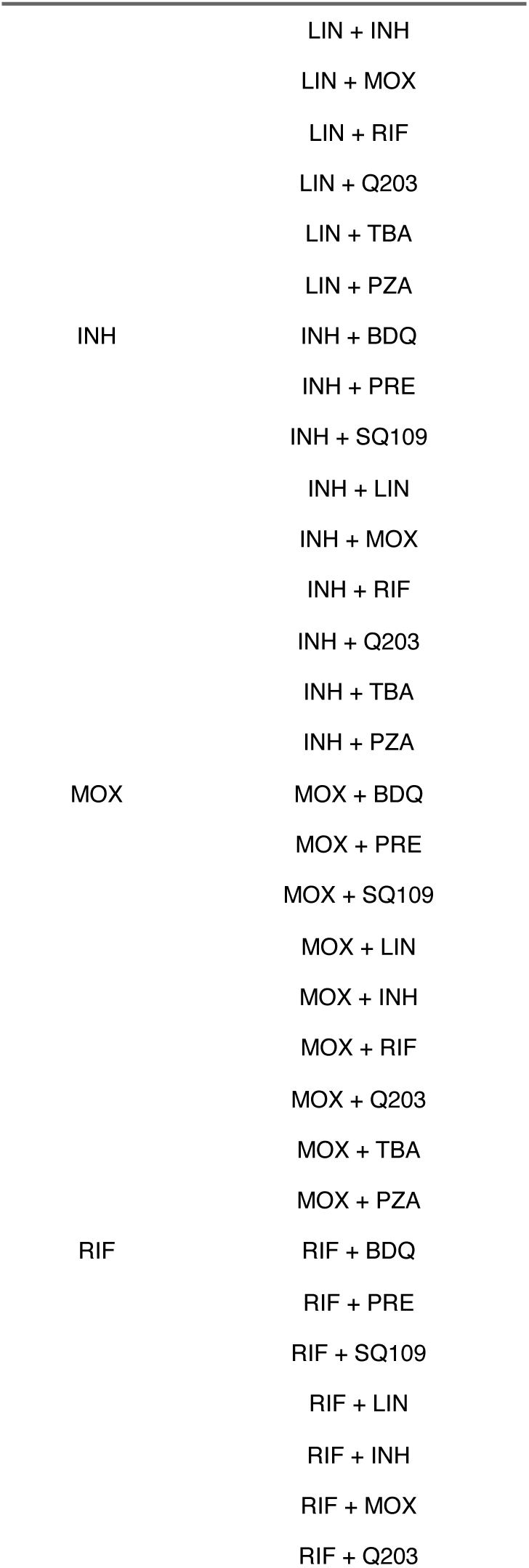
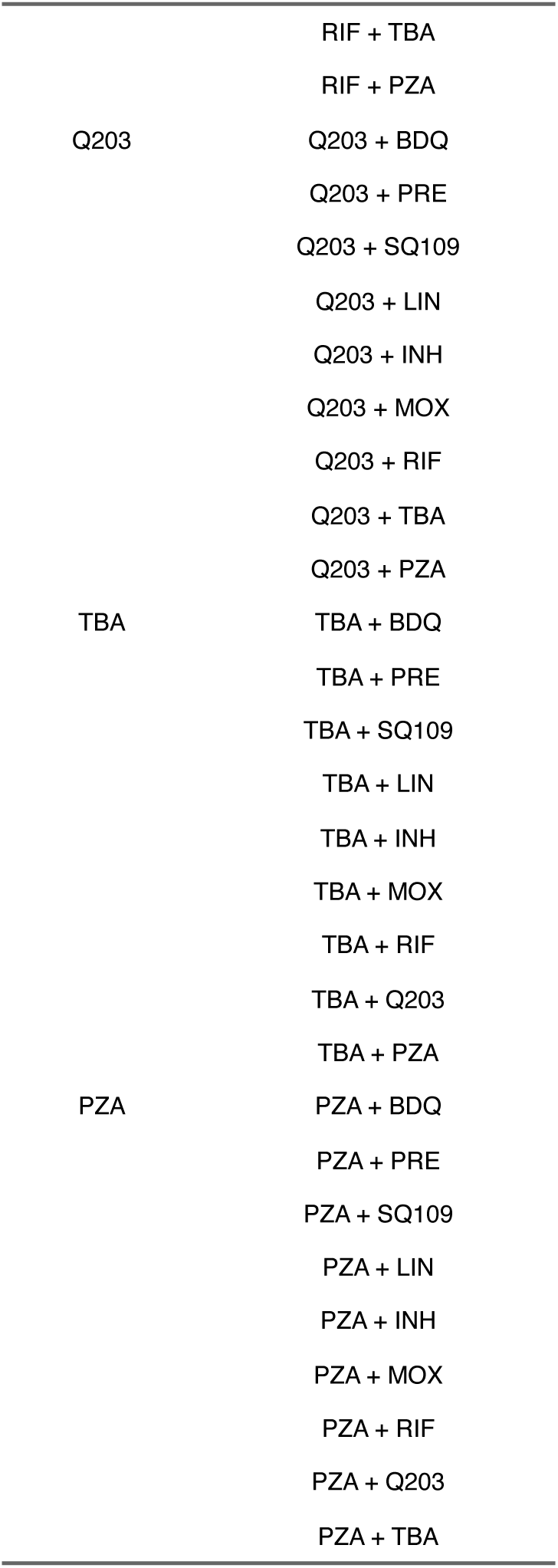
Corresponding drug pairs of each individual drug used in this study. All ten antibiotics and all nine corresponding drug pairs of each individual antibiotic are listed on the left-most and right-most columns, respectively.

**Supplementary Table 3.** All three-way combinations tested in this study.

| Three-way combinations |
| --- |
| BDQ + PRE + LIN |
| BDQ + PRE + INH |
| BDQ + PRE + MOX |
| BDQ + PRE + RIF |
| BDQ + LIN + INH |
| BDQ + LIN + MOX |
| BDQ + LIN + RIF |
| BDQ + INH + MOX |
| BDQ + INH + RIF |
| BDQ + MOX + RIF |
| PRE + LIN + INH |
| PRE + LIN + MOX |
| PRE + LIN + RIF |
| PRE + INH + MOX |
| PRE + INH + RIF |
| PRE + MOX + RIF |
| LIN + INH + MOX |
| LIN + INH + RIF |
| LIN + MOX + RIF |
| INH + MOX + RIF |

**Supplementary Table 4.** Identification and accession numbers of the 14 strains used in this study. For all 14 strains used in this study, the identification name and Sequence Read Archive (SRA) accession numbers are listed in the left and right columns, respectively.

| Strain | SRA accession number |
| --- | --- |
| 044 | SRR5073757 |
| 070 | SRR5074078 |
| 082 | SRR5073981 |
| 084 | SRR5073991 |
| 139 | SRR5073977 |
| 185 | SRR5073510 |
| 245 | SRR5067510 |
| 355 | SRR5067598 |
| 358 | SRR5067503 |
| 414 | SRR5067585 |
| 478 | SRR5065461 |
| 545 | SRR5065319 |
| 617 | SRR5065320 |
| Erdman | SRR24083712 |

**Supplementary Figure 1.**
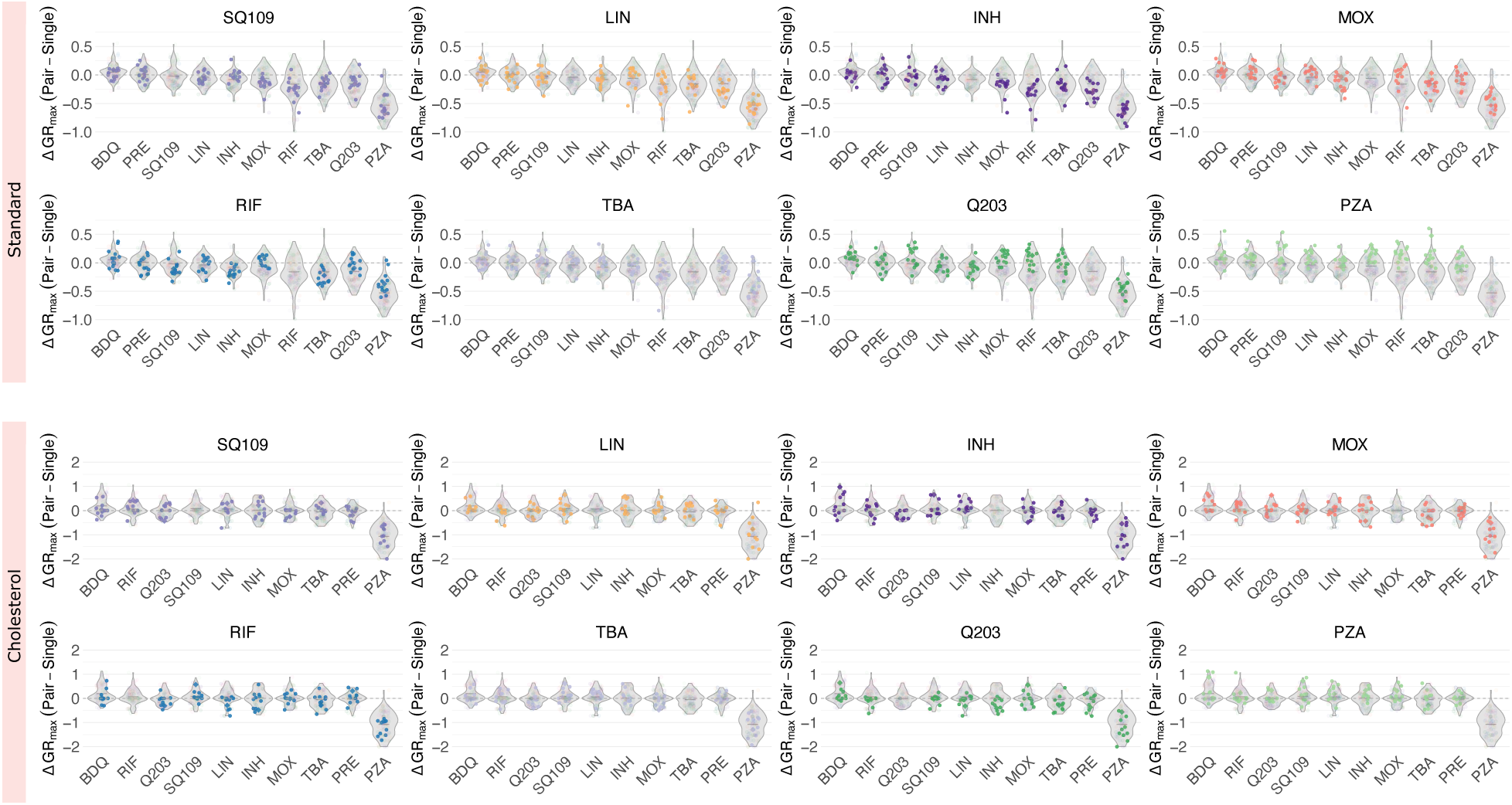
Comparison of GR_max_ values between single drugs and drug pairs. In both standard (top) and cholesterol (bottom) conditions, each of the eight panels represents the change in GR_max_ between a single drug and its 9 corresponding pairs. ΔGR_max_ is calculated by subtracting the GR_max_ value of the single drug from that of a corresponding drug pair. For each single drug, all nine corresponding drug pairs were tested in fourteen clinical isolates in standard and eleven clinical isolates in cholesterol. Each point denotes a GR_max_ value of a drug pair for an individual isolate and represents the median GR_max_ value across three biological replicates. A negative ΔGRmax value indicates an improvement in GRmax when a single drug is combined with another drug.

**Supplementary Figure 2.**
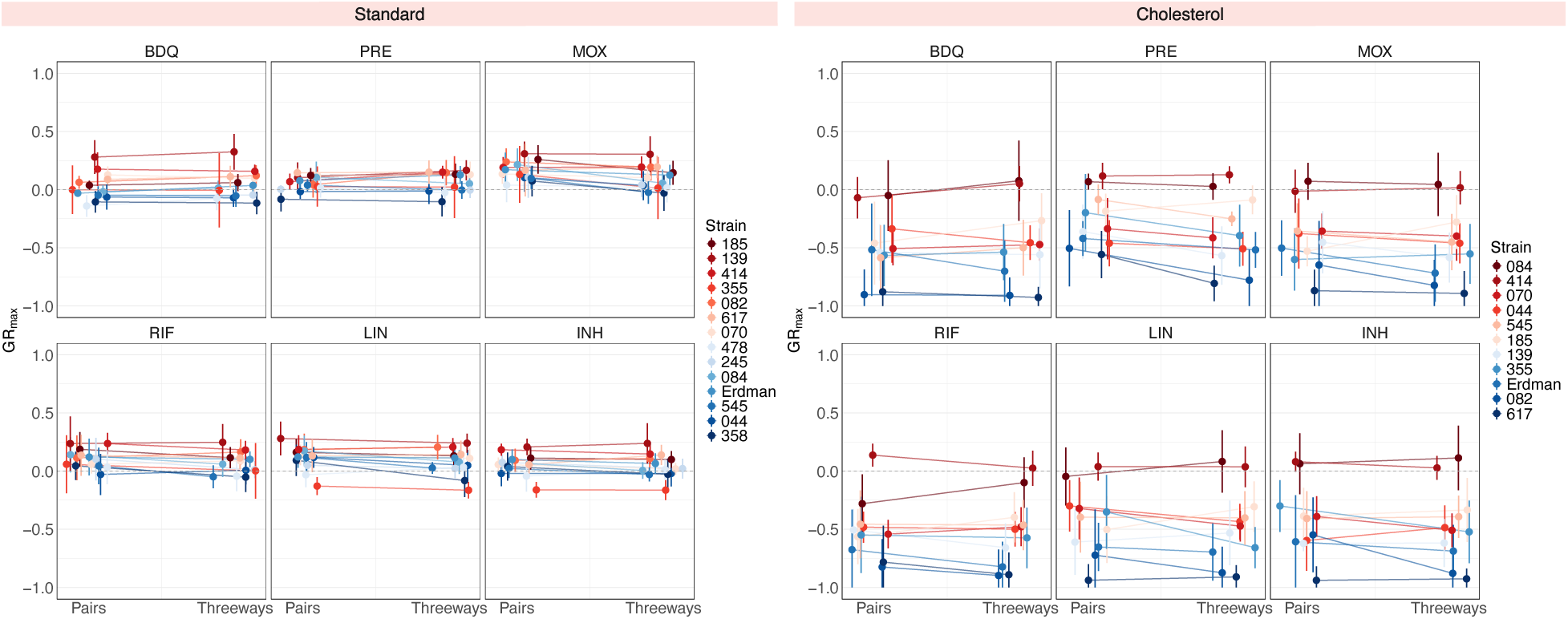
Comparison of GR_max_ values between drug pairs and three-way combinations. In both standard (left) and cholesterol (right) conditions, each of the six panels represents a single drug, with the mean GR_max_ value of its 5 corresponding drug pairs on the left side and the mean GR_max_ value of its ten corresponding three-way combinations on the right side. Vertical bars represent the standard deviation of the mean GR_max_ value. Each point corresponds to one of 14 clinical isolates in standard (left), or one of 11 clinical isolates in cholesterol (right).

**Supplementary Figure 3.**
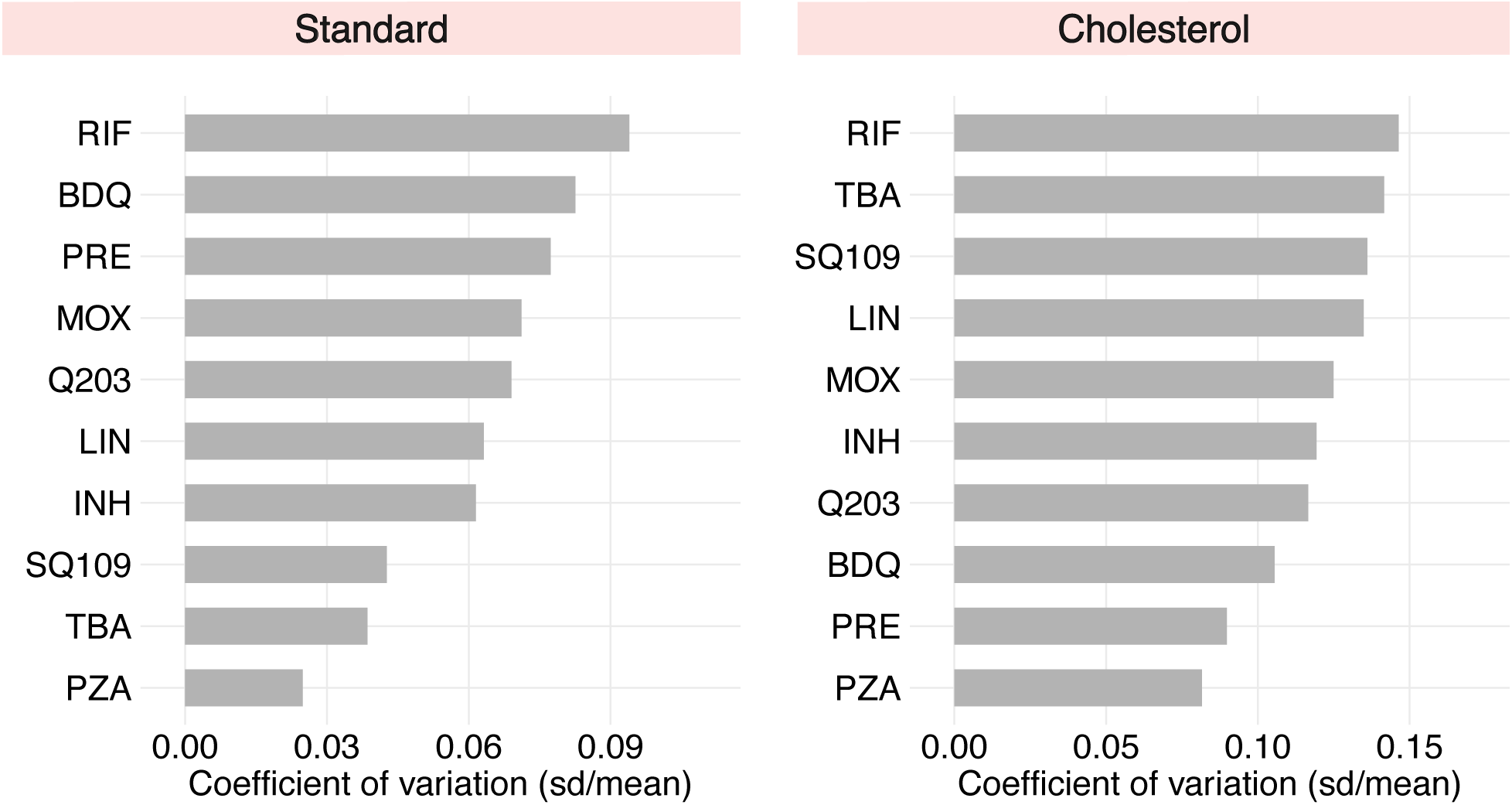
Coefficient of variation (CV) values of ten single drugs. In both standard (left) and cholesterol (right) conditions, the CV was calculated for each drug by dividing the standard deviation of the GR_max_ values by the mean GR_max_ values of 14 isolates in standard and 11 isolates in cholesterol. Drugs were rank ordered from largest to smallest CV (top to bottom, respectively), with larger values indicating greater variation in GR_max_ between strains.

**Supplementary Figure 4.**
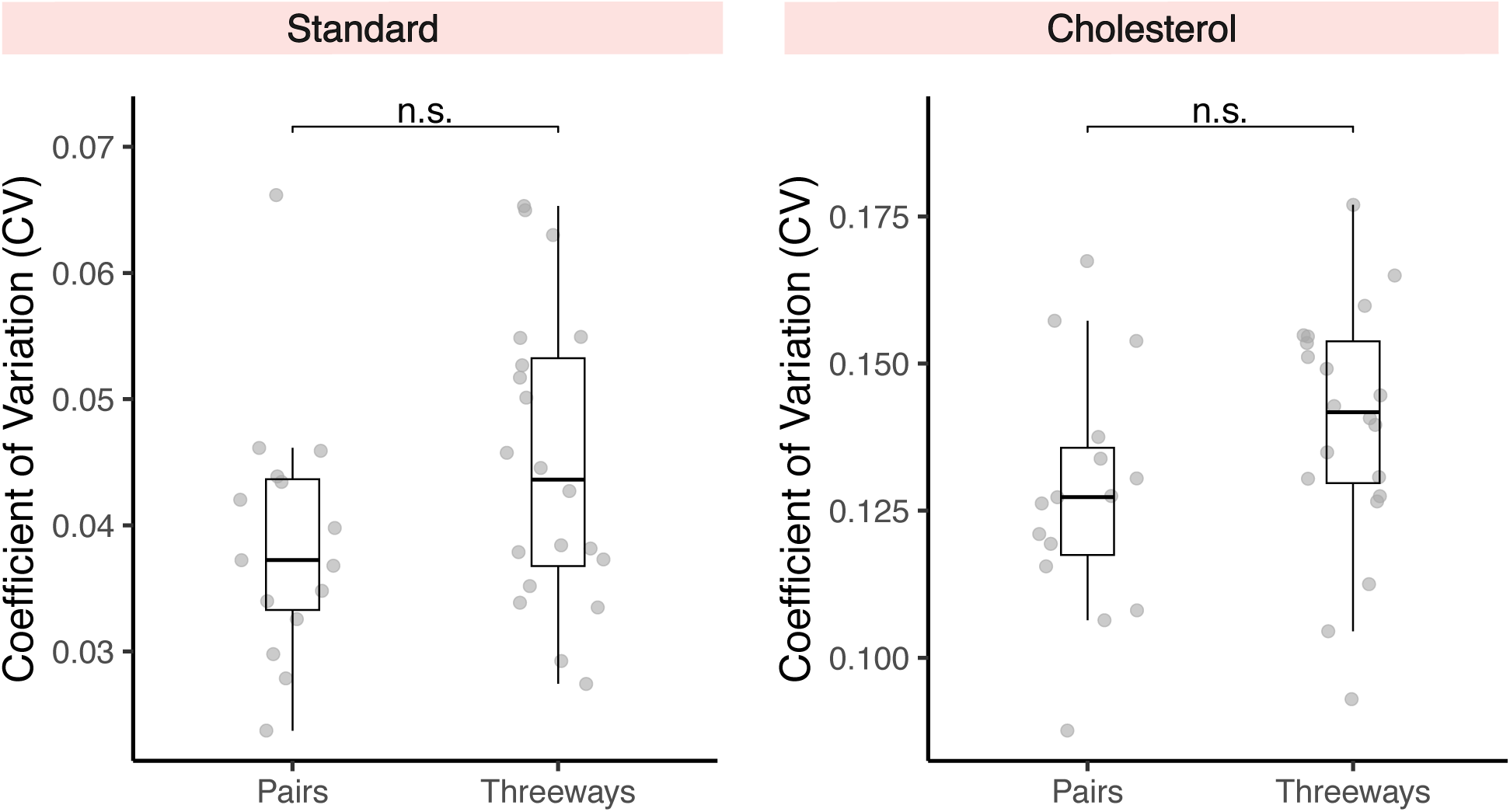
Comparison of mean coefficient of variation (CV) between drug pairs and three-way combinations. In both standard (left) and cholesterol (right) conditions, the CV of each drug pair (total of 15) and three-way combinations (total of 20) was calculated by dividing the standard deviation of the GR_max_ values by the mean GR_max_ values of 14 isolates in standard and 11 isolates in cholesterol. A Student’s t-test was performed to determine whether the difference between the mean CV of drug pairs and the mean CV of three-way combinations is statistically significant.

